# Excess interictal activity marks seizure prone cortical areas and mice in a genetic epilepsy model

**DOI:** 10.1101/2021.12.23.473545

**Authors:** William F. Tobin, Matthew C. Weston

## Abstract

Genetic epilepsies are often caused by variants in widely expressed genes, potentially impacting numerous brain regions and functions. For instance, gain-of-function (GOF) variants in the widely expressed Na^+^-activated K^+^ channel gene *KCNT1* alter basic neurophysiological and synaptic properties of cortical neurons, leading to developmental epileptic encephalopathy. Yet, aside from causing seizures, little is known about how such variants reshape interictal brain activity, and how this relates to epileptic activity and other disease symptoms.

To address this knowledge gap, we monitored neural activity across the dorsal cortex in a mouse model of human KCNT1-related epilepsy using *in vivo*, awake widefield Ca^2+^ imaging. We observed 52 spontaneous seizures and 1700 interictal epileptiform discharges (IEDs) in homozygous mutant (*Kcnt1^m/m^*) mice, allowing us to map their appearance and spread at high spatial resolution. Outside of seizures and IEDs, we detected ~46,000 events, representing interictal cortical activity, in both *Kcnt1^m/m^* and wild-type (WT) mice, and we classified them according to their spatial profiles.

Spontaneous seizures and IEDs emerged within a consistent set of susceptible cortical areas, and seizures propagated both contiguously and non-contiguously within these areas in a manner influenced, but not fully determined, by underlying synaptic connectivity. Seizure emergence was predicted by a progressive concentration of total cortical activity within the impending seizure emergence zone. Outside of seizures and IEDs, similar events were detected in WT and *Kcnt1^m/m^* mice, suggesting that the spatial structure of interictal activity was largely preserved. Several features of these events, however, were altered in *Kcnt1^m/m^* mice. Most event types were briefer, and their intensity more variable, across *Kcnt1^m/m^* mice; mice showing more intense activity spent more time in seizure. Furthermore, the rate of events whose spatial profile overlapped with where seizures and IEDs emerged was increased in *Kcnt1^m/m^* mice.

Taken together, these results demonstrate that an epilepsy-causing K^+^ channel variant broadly alters physiology. Yet, outside of seizures and IEDs, it acts not to produce novel types of cortical activity, but rather to modulate its amount. The areas where seizures and IEDs emerge show excessively frequent and intense interictal activity and the mean intensity of an individual’s cortical activity predicts its seizure burden. These findings provide critical guidance for targeting future research and therapy development.

## Introduction

Many neurological disorders are defined by the symptoms they produce, while the ways in which brain activity is disrupted in the disease state remain mysterious. This is a particularly significant concern for genetic epilepsies, which are often caused by variants in genes that encode common neuronal components, such as ion channels, synaptic proteins, and cell signaling molecules^2^. Because the neurophysiological role of these molecules depends on factors such as cell-type, circuit structure, and developmental stage, the effects of pathological variants are potentially varied and widespread. ^3^ Yet, beyond the uniting fact that they cause seizures and interictal spikes in patients, little is known about how and where they reshape brain activity, and how these actions relate to disease symptoms. This is a critical knowledge gap for two major reasons: (1) the subclinical effects of epileptogenic variants on neural activity may directly contribute to the occurrence of seizures and interictal spikes, and (2) many epileptic patients suffer from neurological co-morbidities that likely result from effects of the variant on brain activity outside of seizure epochs. In both scenarios, these effects represent potentially important therapeutic targets.

The *KCNT1* gene encodes a high conductance Na^+^-activated K^+^ channel, variants in which cause a spectrum of childhood epilepsies and neurodevelopmental disorders.^4–9^ The KCNT1 channel is widely expressed in the brain and mediates a large, delayed, non-inactivating, outward current that is engaged by Na^+^ influx and depolarization.^10–14^ Depending on the voltage range over which the channel is active, its physiological role can span setting the resting membrane potential to contributing to action potential repolarization to mediating slow afterhyperpolarizations following burst firing. Interestingly, epilepsy-causing *KCNT1* variants are almost exclusively gain-of-function (GOF), resulting in an increased current magnitude. In humans, the Y796H variant causes sleep-related hypermotor epilepsy (SHE, formerly autosomal dominant nocturnal frontal lobe epilepsy, ADFNLE).^6,15,16^ Aside from nocturnal seizures, patients carrying this variant exhibit a range of symptom severity from nonambulatory and noncommunicative to a normal neurological exam. Previously, we generated a mouse model in which the mouse ortholog of the human Y796H variant (Y777H) replaced the WT allele. ^17^ Mice homozygous for the YH allele showed reduced excitability in cortical GABAergic neurons, increased homotypic synaptic connectivity, and frequent spontaneous seizures, but normal lifespan and general health, making it an ideal model in which to monitor seizures and IEDs alongside interictal cortical activity.

Macroscopic widefield calcium imaging allows measurement of bulk neural activity at centimeter scale with high spatial resolution (~25μm) across the dorsal cortex of awake and behaving mice.^18^ It has been used in healthy mice to identify correlates of sensory stimulation, movement, numerous behavioral task-related variables, and learning and memory.^19–25^ It is also well-suited to the study of epileptic pathology, which often involves brain-wide networks of connected areas.^26^ Previous studies have imaged pharmacologically induced focal seizures and IEDs in the cortices of WT rodents ^27–30^, and epileptic activity caused by glioma.^31,32^ These studies have concluded that induced epileptic activity spreads within synaptic-coupled areas, and to areas where inhibition is compromised.

Despite the elegance of this work, pharmacologically- or glioma-induced seizures may differ significantly from spontaneous seizures in a genetic model, and the effects of disease-causing gene variants on neural activity at this scale are relatively unexplored. Here, we used macroscopic widefield calcium imaging to map spontaneous seizures and IEDs in the *Kcnt1^m/m^* cortex. These maps provide a detailed description of pathological activity in the chronically epileptic brain and can be used to target future experiments at the cellular scale to regions with an established link to disease. Furthermore, comparing interictal activity in *Kcnt1^m/m^* and WT mice allowed us to identify how the YH variant altered neural activity outside of seizures and IEDs. We discovered specific associations between alterations in interictal activity and susceptibility to seizures and IEDs; relationships that provide new insights into the features of neural activity that underlie epileptic pathology.

## Materials and Methods

### Mice

The animals included in this study were housed and used in compliance with the National Institutes of Health Guidelines for the Care and Use of Laboratory Animals and approved by the Institutional Animal Care and Use Committee at the University of Vermont. Experimental mice were generated by first mating the *Kcnt1*^Y777H^ knockin line ^17^ to the *Snap25*-GCaMP6s line (Jackson Labs Stock No: 025111).^33^ Double heterozygous offspring were then mated to generate mice carrying the Snap25-GCaMP6s construct that were either homozygous Y777H or WT at the *Kcnt1* locus. *Kcnt1*^Y777H^ genotyping was performed as previously reported^17^ and Snap25-GCaMP6s was genotyped following the protocol supplied by Jackson Labs.

### Surgery

All surgical procedures were performed on mice under 1-2% isofluorane anesthesia. The mouse’s eyes were coated with protective ophthalmic ointment (Paralube^®^, Dechra Pharmaceuticals) and scalp hair removed, before fixing the head in a stereotaxic frame (Stoelting, Mouse and Neonates Adaptor, Item # 51625) and placing a homeothermically controlled heating pad under the body (PhysioSuite, Kent Scientific, item # PS-02). Prior to removing the scalp skin with fine surgical scissors (FST Item No. 14058-09), it was scrubbed with 10% povidone-iodine followed by 70% ethanol, three times. Once the skull was exposed and overlying fascia removed, we retracted neck muscles, bilaterally, at their anterior most attachment point on the interparietal bone. We glued the marginal, cut skin in place with a tissue adhesive (3M Vetbond). For one animal, the dorsal cranium was replaced with a large glass window (Crystal Skull, Labmaker) following the protocol detailed in Kim et al..^34^ Next, an aluminum head plate, fabricated following the design in Goldey et al.^35^, was fixed to the skull using UV curing cyanoacrylate (Loctite 4305). To minimize the escape of excitation light from the implant during imaging, powdered black tempra paint (Jack Richeson & Co SKU# 101508) was mixed into the glue before curing. For transcranial imaging, we applied a thin layer of UV curing cyanoacrylate to the surface of the skull. After surgery, the head plate was filled with silicone elastomer (Kwik-Sil, WPI) for protection during recovery. We treated postoperative pain with dual antibiotic and anesthetic cream (Neosporin Plus) and ketoprofen (5 mg/kg i.p.) and waited at least 24 hr before imaging.

### Data Acquisition

We used a custom tandem-lens epifluorescent macroscope ^36^ with 50- and 105-mm focal length objective and image forming lenses, respectively (Nikon Nikkor 50mm f/1.4 and 105mm f/1.8), for 2.1X magnification. The light source was a broadband LED (X-Cite 120 LED) with a GFP filter set (Chroma 49002) in 50-mm circular glass, mounted in Thorlabs SM2 hardware in the configuration illustrated in Fig. 1A. In several sessions collected early during this study (5.4 hr of imaging collected over 9 days from 1 mouse and containing 18 seizures) only epifluorescence illumination was used. However, in all other sessions, we used a 530-nm LED (Thorlabs, M530F2, LEDD1B) coupled to an optical fiber pointed obliquely at the cortical surface to illuminate every other frame. The resulting reflectance images were used to correct the GCaMP signal for hemodynamic artifacts. We included seizure data from sessions lacking reflectance images in Fig.1-4 but excluded these sessions from analyses in Fig. 5-7 where a fair comparison of event duration and intensity was not possible. To toggle illumination, both LEDs were driven using the camera frame exposure signals and routed with a microcontroller board (Arduino Uno). All images were captured with an Andor Zyla 5.5 sCMOS camera (Camera Link 10-tap) controlled by the open-source microscopy software μManager^37^ (version 1.4) from within Matlab.

**Figure 1.**
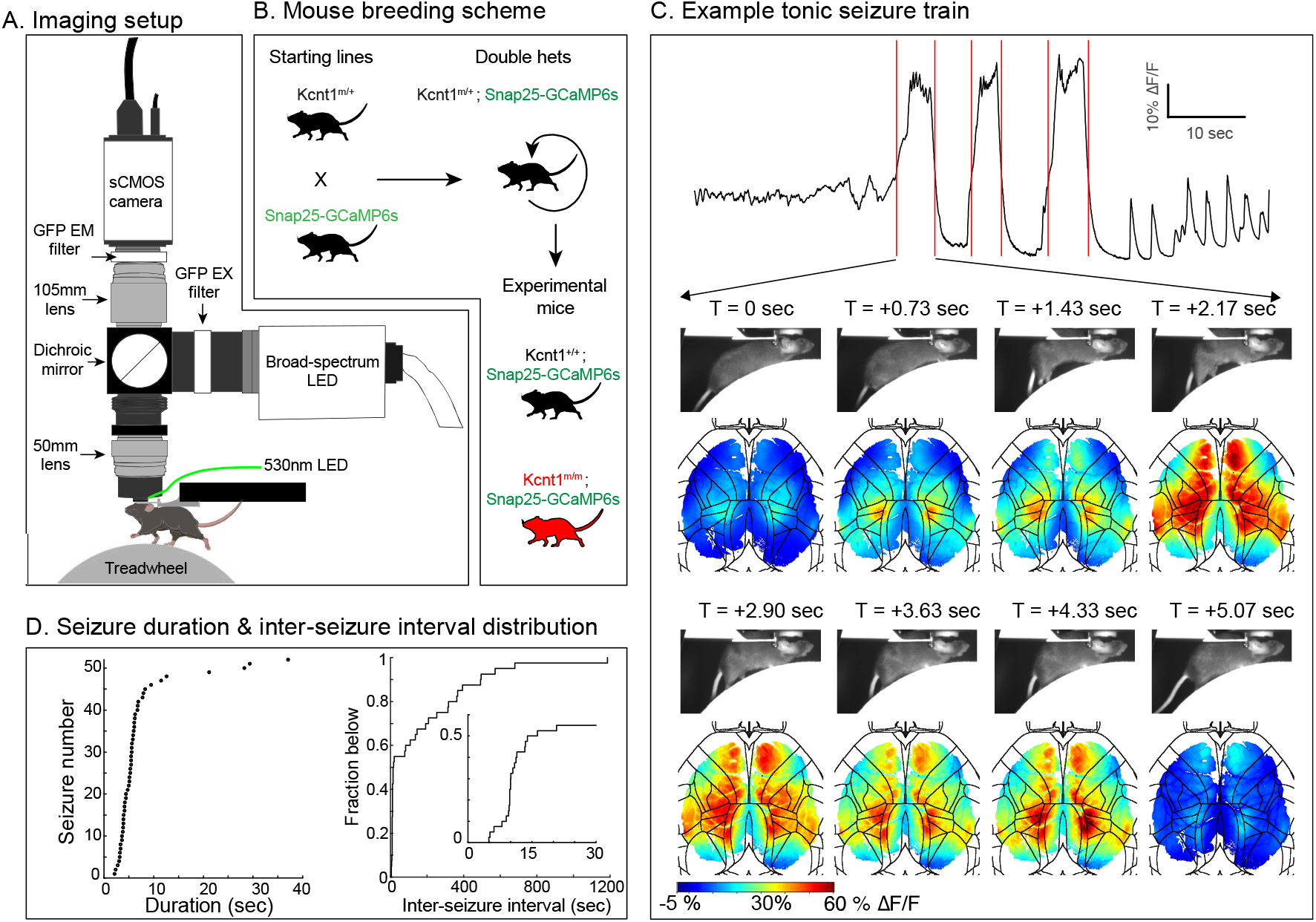
Widefield imaging of dorsal cortex in *Kcnt1^m/m^* and WT mice. (**A**) A schematic of the tandem-lens, epifluorescent macroscope used for *in vivo*, awake widefield Ca^2+^ imaging. Every other frame was illuminated with a 530 nm LED pointed obliquely at the skull surface; the resulting reflectance images were used to correct our GCaMP6s signal for hemodynamic artifacts. (**B**) The breeding scheme used to generate experimental mice. (**C**) An example tonic seizure train. The trace shows average ΔF/F signal calculated across all pixels involved in at least one of the three seizures shown. Red lines demarcate seizure boundaries. Images below show still frames of mouse body video and simultaneous ΔF/F frames of the dorsal cortex, with black lines marking the Allen Mouse CCF area borders. Time relative to seizure start is indicated above each pair of frames. (**D**) Most seizures were brief and clustered in time. The scatter plot on the left shows the duration of all seizures. The plot on the right displays an empirical cumulative distribution function of inter-seizure interval generated from all sessions containing more than one seizure. The inset shows a zoom to the foot of the plot.

**Figure 2.**
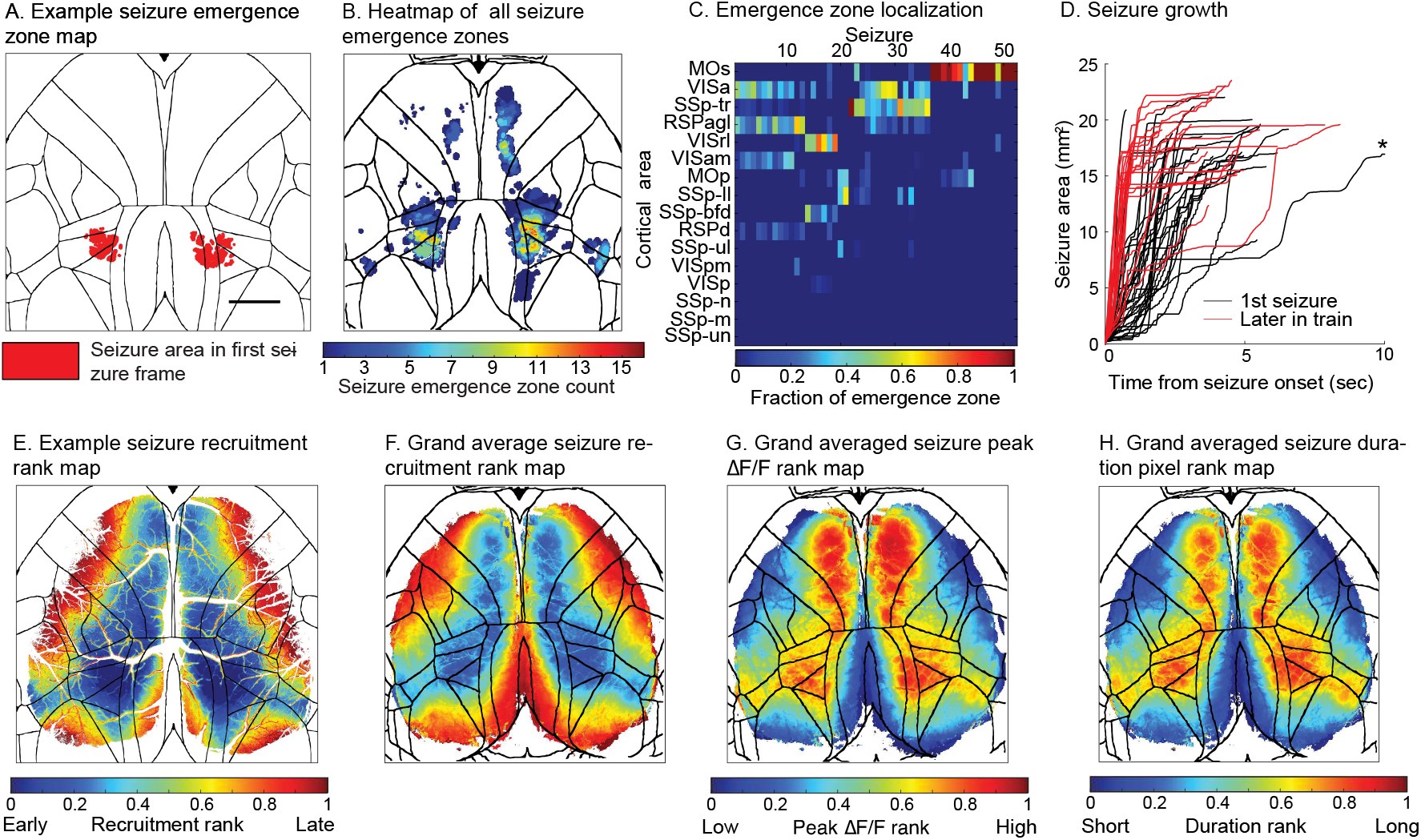
Spontaneous seizure mapping identifies seizure susceptible cortical areas in *Kcnt1^m/m^* mice. (**A**) An example seizure emergence zone map. Red pixels are those above seizure threshold in the first frame containing seizure activity. Black lines mark the borders of the Allen Mouse CCF areas. Scale bar is 1 mm. (**B**) A heatmap of seizure emergence zones summed over all 52 seizures. The value of each pixel indicates the number of seizures in which that pixel was a part of the emergence zone. (**C**) A heatmap showing the fraction of each emergence zone located within each of the 16 CCF areas imaged in all sessions. (**D**) A plot showing the area of seizing tissue as a function of time, covering the period between the first seizure frame and maximal seizure area. Black traces represent the first seizure in seizure trains or seizures that occurred singly, and red traces the seizures that occurred after the first in seizure trains. Asterisk marks a single trace that extended beyond the limit of the x-axis. (**E**) An example seizure recruitment rank heatmap (same seizure as panel A). The value of each pixel is its recruitment time rank. (**F**) A grand average seizure recruitment rank heatmap, taken across all individual mouse average maps after co-registration. (**G**) A grand average seizure peak ΔF/F rank heatmap, taken across individual mouse averages. (**H**) A grand average heatmap of ranked seizure duration at each pixel, taken across individual mouse averages.

**Figure 3.**
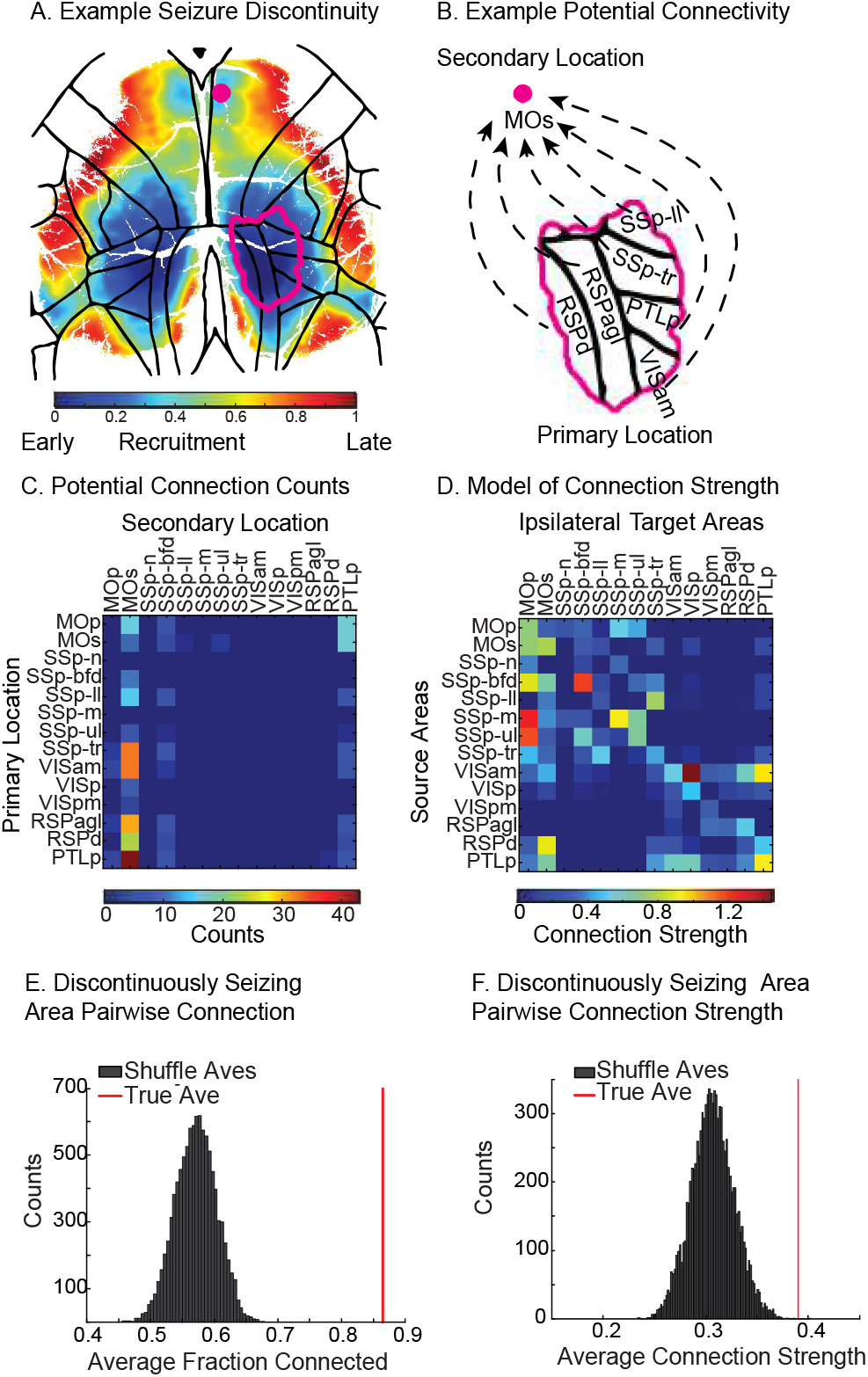
Normal cortico-cortical synaptic connectivity predicts long range seizure jumps in *Kcnt1^m/m^* mice. (**A**) An example seizure rank recruitment heatmap showing spatial discontinuity in seizure propagation. The magenta border indicates the extent of seizing tissue in the primary location at the time the seizure emerged at the secondary location, marked by a magenta dot in MOs. (**B**) A diagram indicating potential synaptic connections that are hypothesized to mediate propagation between each of the areas in the primary and secondary seizure locations. (**C**) A heatmap indicating the number of times each pair of cortical areas was found in primary and secondary seizure locations. (**D**) A heatmap showing area-to-area connection strength in the inter-area connectivity model adapted from Oh *et al*.^1^ cropped to include only areas in our FOV (**E**) Results of a simulation to test primary-to-secondary area connection probability against a random draw from the model in D. In each repetition (*n* = 10,000), the primary location was rotated and repositioned and a secondary location (>300μm distant) was selected at random within a hemisphere. To simulate bilaterally symmetrical seizures, we applied the same position change and used the same secondary location in both hemispheres. We calculated the fraction of connected areas in each simulated discontinuity and the grand average across all discontinuities, simulated grand averages are shown by the histogram and the true grand average by the red line. (**F**) Results of a shuffle analysis to compare the connection strength between primary and secondary seizure locations to a random draw from the connectivity model in D. In each repetition (*n* = 10,000), all non-zeros connection weights in the true primary-to-secondary data set were replaced with connection weights drawn randomly from the model. For bilaterally symmetrical seizures, we used the same random value for equivalent area pairs in both hemispheres. We calculated the average connection strength for each simulated discontinuity and then the grand average across all discontinuities. Histogram shows simulated grand averages, and the red line marks the true grand average connection weight.

**Figure 4.**
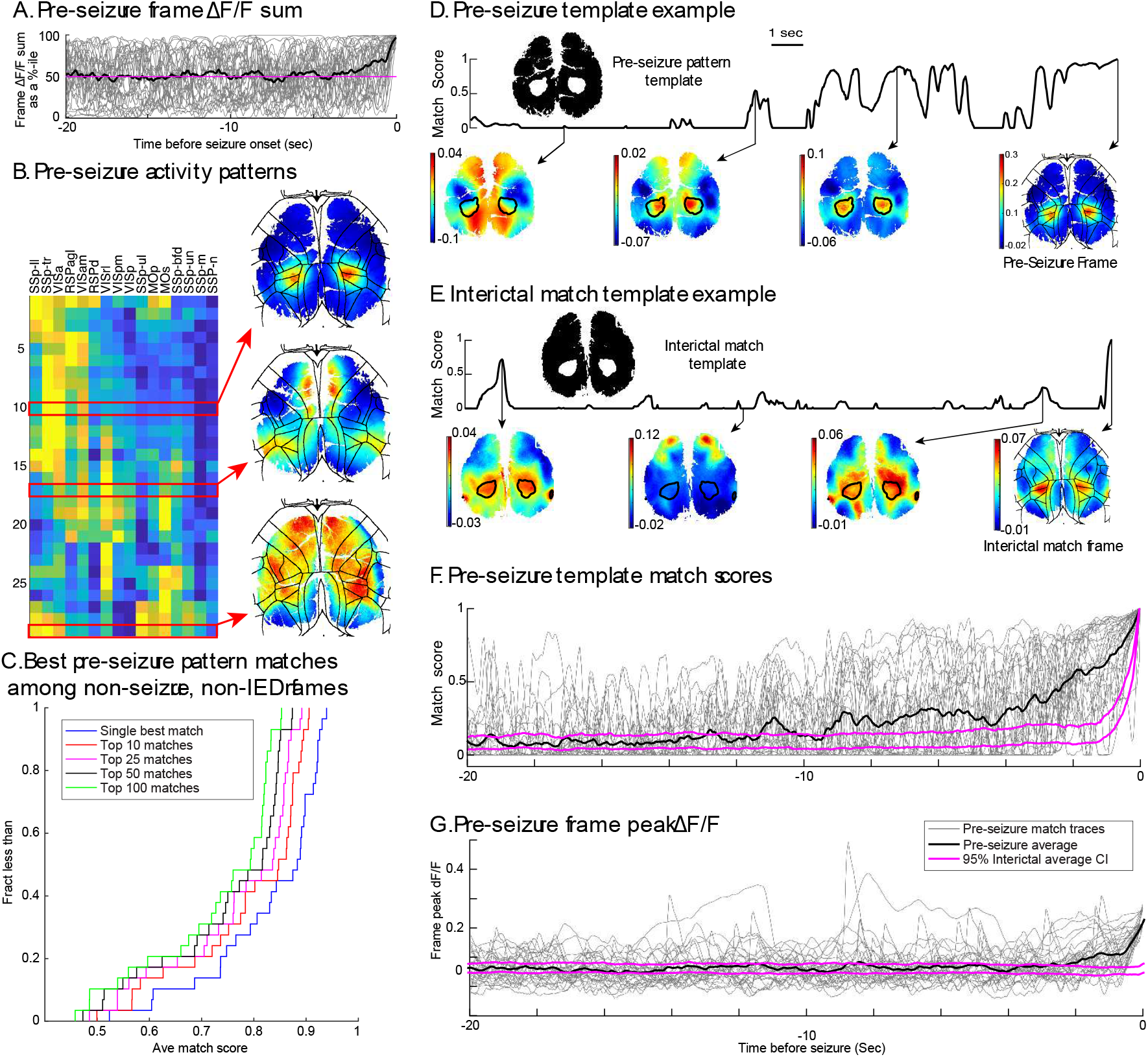
Abnormally persistent activity in the seizure emergence zone precedes *Kcnt1^m/m^* seizure onset. (**A**) Summed ΔF/F across all pixels covering cortical tissue in the lead up to seizure as a percentile of all frame sums during interictal activity. Individual traces in grey, average across all 29 seizures preceded by at least 20 sec of seizure-free activity in black. Magenta line marks 50^th^ percentile. Percentiles calculated separately for data collected without hemodynamic correction (*n* = 8 seizures) (**B**) A heatmap showing Allen Mouse CCF area average ΔF/F values in the last frame before emergence of all 29 seizures in our data set preceded by at least 20 sec of seizure-free activity. Rows are ordered according to optimal leaf order after hierarchical clustering. Three example pre-seizure frames are shown on the right with CCF area boundaries overlaid. (**C**) A plot showing the empirical cumulative distribution functions of single best match scores and average match scores across top 10, 25, 50, and 100 matches to each pre-seizure activity patterns. (**D**) An example pre-seizure template match score trace in the 20 sec lead up to seizure onset. The pre-seizure template, derived from the pre-seizure frame (far right) identifies pixels containing peak activity (highest 5% of ΔF/F values) in the last frame before seizure onset. Example frames, with template boundaries overlaid, are shown for three points along the trace. (**E**) An example interictal template match score trace in the 20 sec lead up to one of the top 50 interictal matches to the pre-seizure activity pattern shown in panel D. The interictal match template, derived from the interictal match frame (far right), was used to generate the match score trace. Example frames, with template boundaries overlaid, are shown for three points along the trace. (**F**) A comparison of template match scores in the 20 sec lead up to seizures, and interictal match frames. Gray traces show individual pre-seizure template match scores, and the black trace shows the average across all pre-seizure traces. Magenta lines show the 95% confidence interval for a match score average calculated over a random selection from the top 50 interictal matches to each pre-seizure template. (**G**) A comparison of frame peak ΔF/F in the lead up to seizures, and interictal match frames. Gray traces show frame peak ΔF/F in the lead up to each seizure, and the black trace is the average across all pre-seizure traces. Magenta lines show the 95% confidence interval for a frame peak ΔF/F average calculated over a random selection from the top 50 interictal matches to each pre-seizure template.

**Figure 5.**
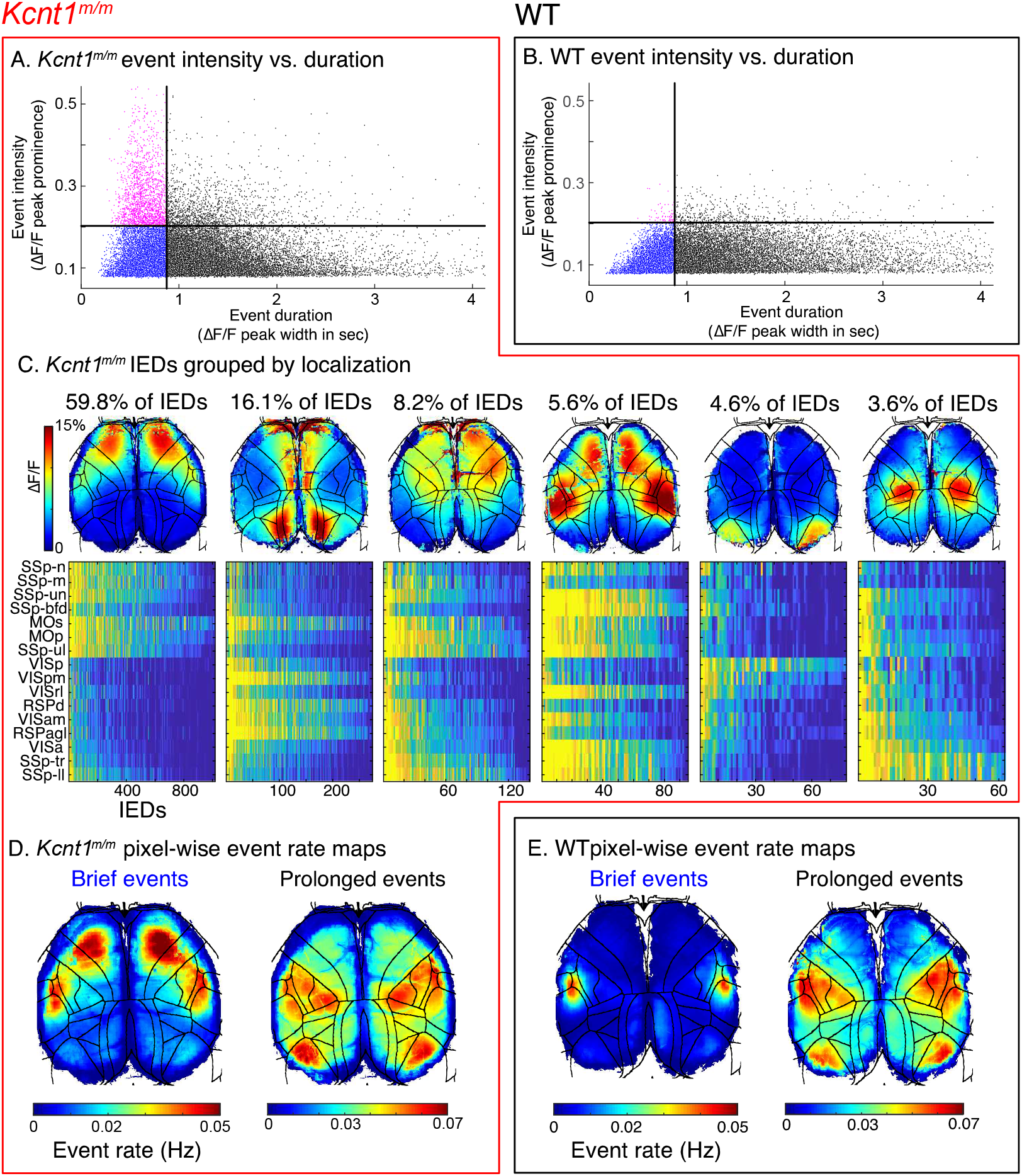
IEDs constitute the high intensity tail of excessive activity centered in *Kcnt1^m/m^* MOs. (**A** and **B**) Scatter plots of event intensity versus duration for events detected in *Kcnt1^m/m^* (**A**) and WT (**B**) mice. Event intensity was measured as the 99^th^ percentile of ΔF/F peak prominences on all event pixels and duration was measured as the mean with at half peak prominence calculated over all event pixels. Horizontal lines mark IED intensity threshold (0.20), and vertical lines mark IED duration threshold (0.875 sec). Plots were cropped to highlight region relevant to IED detection. (**C**) Images show the average ΔF/F frame taken across all the IEDs within clusters produced by hierarchical clustering. The value of pixels outside the IED were set to zero in frames contributing to these averages. Above each image, the percentage of all *Kcnt1^m/m^* IEDs contained in the cluster is given. Three small clusters collectively representing 1.9% of IEDs are not shown. Below, individual heatmaps show average ΔF/F in each cortical area for all IEDs in the cluster. Values in each column are scaled to span 0 to 1. See Experimental Methods for discussion of cluster number selection (**D** and **E**) Pixel-wise rate maps for events that meet IED duration but not intensity criteria (‘Brief Events’) and all other events (‘Prolonged Events’) in *Kcnt1^m/m^* (**D**) and WT (**E**) mice. Maps are grand averages taken across individual mouse session averaged rate maps.

**Figure 6.**
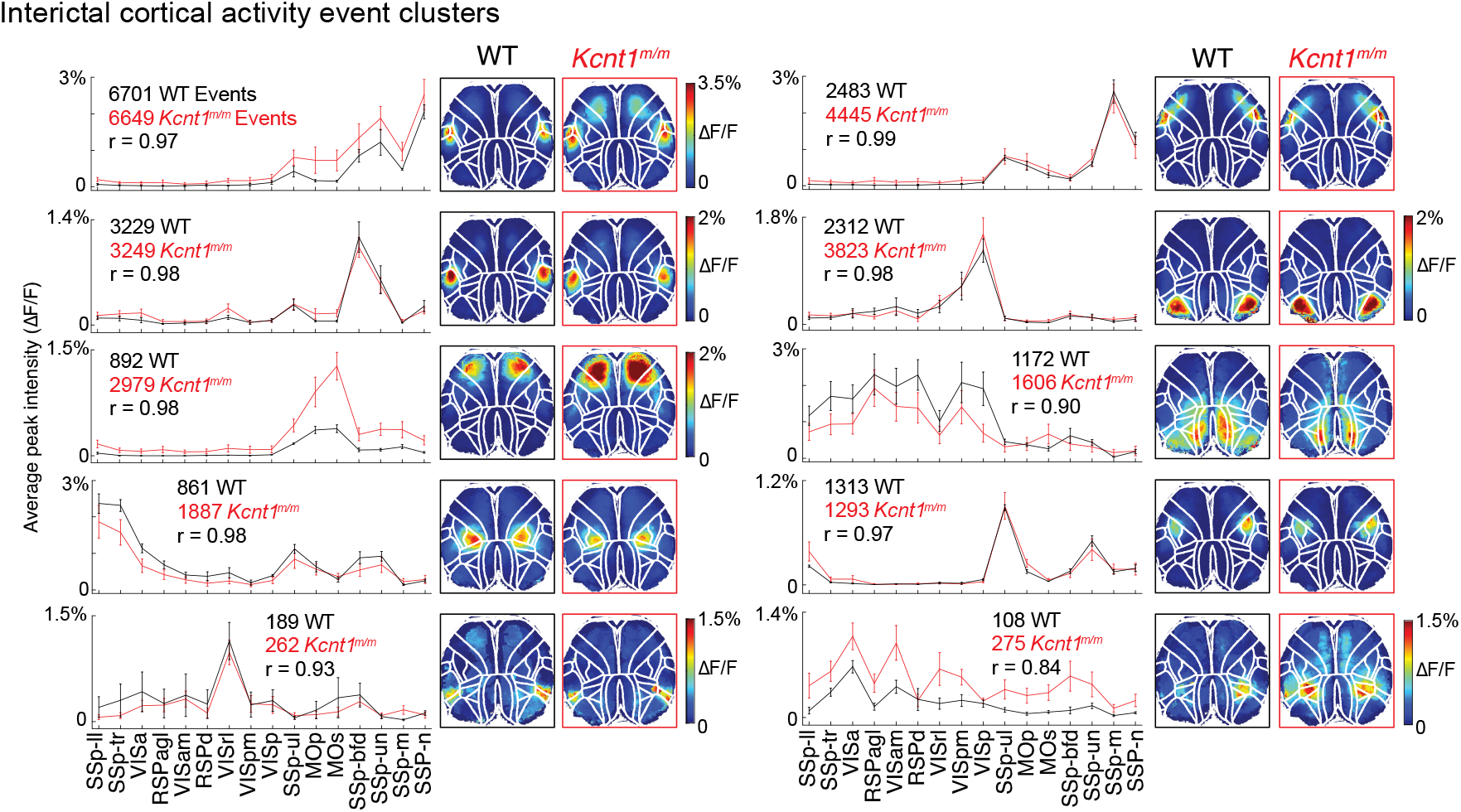
Spatial organization of interictal cortical activity is largely preserved in *Kcnt1^m/m^* mice. Line plots show the average peak intensity of neural activity within each Allen Mouse CCF area we imaged during events in each cluster. These values were constructed by taking the average ΔF/F peak value across all pixels involved in an event in each area, pixels not involved in the event were set to zero. Lines represent grand averages for each group, taken across session averages for each mouse. Error bars show SEM across mouse averages. Text above each plot gives the total number of events in the cluster from each group and the *Kcnt1^m/m^* to WT Pearsons correlation coefficient. Images to the right of each line plot show the average event peak intensity at all pixels for each genotype and cluster. These were calculated using event frames in which each pixel was assigned its ΔF/F peak prominence value, and pixels not involved in the event were set to zero. The color bar in the top left applies to all images in the figure except for those where another color bar is present.

**Figure 7.**
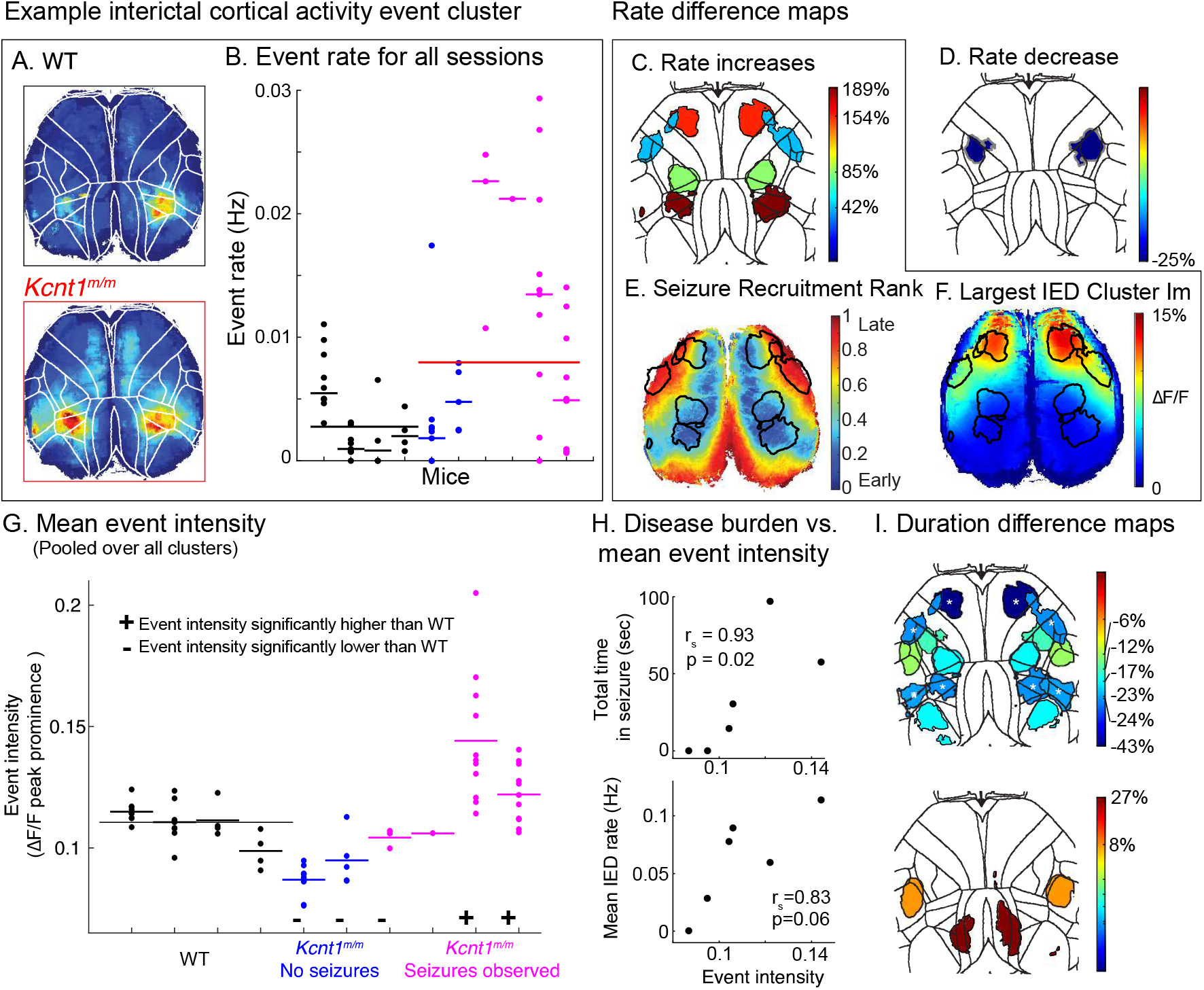
Excessive interictal activity in *Kcnt1^m/m^* cortical areas and mice prone to seizure and IED. (A) Example cluster event peak intensity images from WT and *Kcnt1^m/m^* mice (B) Scatter plot of example cluster event rate for all sessions in each mouse in our data set. Short lines represent animal mean rates and long lines indicated group means weighted by session number. Black indicates WT, blue and magenta points *Kcnt1^m/m^* mice following the same sorting and color scheme as in panel G, the long red line shows the *Kcnt1^m/m^* weighted group mean. (C & D) Heatmaps indicating clusters in which event rate is significantly increased or decreased, respectively, in *Kcnt1^m/m^* mice (Permutation Test, p<0.05 Bonferroni-Holm corrected). Patch color indicates the fractional change in event rate and the borders demarcate peak activity in the cluster event mean image. (E & F) To highlight the correspondence between seizure and IED susceptibility and an increased rate of interictal activity, we have reproduced the grand average seizure recruitment rank map from Fig. 2F and the average image for the largest IED cluster shown in Fig. 5C and drawn on peak activity borders for clusters with increased event rates in *Kcnt1^m/m^* mice. (G) Scatter plot of session mean event intensities (see Experimental Methods). Short lines indicate mouse means with WT in black and *Kcnt1^m/m^* in blue and magenta. *Kcnt1^m/m^* mice are sorted in ascending order, from left to right, by the total seizure time observed in each individual during the sessions represented in this plot. Whether each *Kcnt1^m/m^* mouse differed from WT in terms of their mean event intensity is marked with the + and - symbols described in the legend (Permutation Test, *p* < 0.05 Bonferroni-Holm corrected). (H) Scatter plots of *Kcnt1^m/m^* mouse mean event intensity (short lines in (G)) versus total time in seizure and mean IED rate. Spearman’s correlation coefficient and associated p values are given. (I) Heatmaps indicating clusters in which event duration is significantly increased or decreased in *Kcnt1^m/m^* mice (Permutation Test, *p* < 0.05 Bonferroni-Holm corrected). Patch color indicates the fractional change in event duration and the borders demarcate peak activity in the cluster event mean image.

Mice were secured under the objective lens using thumb screws on fixed threaded rods inserted through the outermost holes of their head plate. The rods were mounted on custom steel plates attached to 25-mm square optical rails (Thorlabs, 9 in long), supported by post bracket platforms (Thorlabs, C1515) on 1.5” optical posts (Thorlabs, P14). Mice were supported a treadwheel made from a Styrofoam cylinder (Golda’s Kitchen, 8” x 4” cake dummy) mounted on a custom axle; wheel position was recorded with an optical shaft encoder (US Digital S6-2500-236-IE-S-B) and a PCIe DAQ system (National Instruments, Part #s 781045-01, 782536-01, 192061-01). Video of the mouse’s body was collected under infrared illumination (Camera: Allied Vision GC750).

### Image Processing

All data processing was performed using Matlab, either on the acquisition PC (Thinkmate VSX R5 540V3) or on the Vermont Advanced Computing Center. Raw images were saved as TIFF files. We subtracted a dark frame, exposed at the start of each session with no illumination, from all frames before de-interleaving reflectance and epifluorescence images. We masked out all pixels not covering the brain, as well as the largest visible blood vessels, using a brightness threshold before transforming GCaMP6s fluorescence values into ΔF/F on each pixel, using the 20^th^ percentile value as an estimate of baseline fluorescence. Similarly, for sessions in which reflectance images were captured, we calculated fractional reflectance on each pixel as (Ref-Ref_o_)/Ref_o_, using the pixels average value as Ref_o_ and subtracted this from GCaMP6s ΔF/F. We then performed singular value decomposition (SVD), using code adapted from Steinmetz & Peters ^38^ and used the first 50 to reconstruct images for all analyses. In three imaging sessions, seizure activity transitioned to cortical spreading depression (CSD); CSD activity was removed before performing SVD because it distorted the resulting singular values.

### Seizure Identification

To identify seizures, we first detected frame spans in which cortical activity was high and changing rapidly using two vectors: (1) the fraction of pixels above 20% ΔF/F in each frame and (2) the average pixel-wise ΔF/F difference between successive frames. After smoothing both with a 10 sec moving average filter, we detected coincident peaks (within 15 sec) in the envelope of vector (1) and vector (2) using the Matlab built-in function findpeaks(). We then screened potential seizures for behavioral symptoms by watching side-by-side videos of the mouse’s body and cortical ΔF/F over the period when vector (1) was above its average value on either side of a peak. In sessions containing 18 seizures, we were unable to align brain and behavior videos and made our judgement based on watching cortical activity videos alone.

For each seizure with behavioral correlates, we next determined the precise borders of seizure activity by using a per-animal ΔF/F threshold set to the 99^th^ percentile of non-seizure, non-IED, interictal cortical activity event peaks (see below). This threshold was determined separately for sessions with only epifluorescence illumination. For each seizure, we recorded the first frame in which at least 1% of pixels were above threshold (seizure start), the first frame in which the number of pixels above threshold was maximal (end of seizure growth phase), and the first trailing frame in which less than 1% of pixels was above threshold (seizure termination). Because IEDs sometimes occurred close to seizures and involved above-threshold activity, any “seizures” that met our IED criteria were eliminated.

For picrotoxin seizures, we modified this method by increasing the findpeaks() threshold used for vector (1) from 5% to 60% and by using vector (1) instead of vector (2) to define the video inspection frame span because it showed these seizure boundaries more clearly. Finally, because picrotoxin IEDs did not always meet our spontaneous IED identification criteria, “seizures” with a duration less than 5 sec were discarded.

### Image Alignment and Registration

A nonreflective similarity transformation based on manually placed control points was used to register images from each session to the Allen Mouse Common Coordinate Framework (CCF) v3. Adapting previously published methods, these points were placed on the anterior edge of the cortex, midway along the mediolateral extent of each olfactory bulb and at the midline, as well as on the posterior edge at the midline.^20,39^ A histogram equalized average GCaMP6s fluorescence image was used as reference to estimate these locations in the FOV.

Images from all sessions were co-registered with a three-step process: (1) a within-animal affine transformation based on histogram equalized session average GCaMP6s images (2) an across animal registration based on the functional topography revealed by seed pixel correlation maps (SPCMs) (code adapted from Steinmetz & Peters ^38^). For each session, we generated SPCMs using a 50-by-50 grid of seed pixels, calculated the average map from seed pixels in primary visual cortex (VISp), and took the grand average across all sessions from a given mouse. We used these to identify a similarity transformation to align images across mice. (3) Finally, the cross-animal registration was refined by calculating another similarity transformation based on each mouse’s grand average VISp SPCM and an average VISp SPCM taken across mice following the second step.

### Seizure discontinuity identification

To identify seizure discontinuities, we used the Matlab built-in function imextendedmin() on seizure recruitment rank maps after pre-processing with the following steps: (1) Set the value of all pixels outside the brain to the average value of brain pixels, (2) Interpolate over gaps corresponding to surface vasculature using the built-in Matlab function regionfill(), and (3) Filter the image with a gaussian smoothing kernel with a std of 4.

### Pre-seizure image template matching

To generate pre-seizure activity templates, we processed each pre-seizure frame using the same 3 steps described in the previous section. The template was a binary mask covering the top 5% of pixel values in the resulting image. We created equivalent templates for every other frame in our data set and calculated a ‘match score’ between each pre-seizure template and every other frame template. This score was the number of overlapping template pixels divided by the total number of pixels in the smaller mask; mask size varied slightly across individuals.

To construct the null distribution of match scores preceding interictal matches used in Fig. 4F and 4G we randomly selected an interictal match to each pre-seizure template from among its top 50 best matches. We then averaged individual match score traces covering the 20 sec before each interictal match frame and repeated this process 500 times to estimate 95% average match score confidence intervals. To control for the fact that pre-seizure match scores come from templates derived from pre-seizure frames, not interictal match frames, we used templates derived from each interictal match frame in constructing our null.

### Event detection and peak intensity measurement

We performed event detection in two steps: (1) For each session, we calculated average ΔF/F within regions of interest (ROIs) covering the FOV in a 50-by-50 grid and detected peaks in these signals using the Matlab built-in function fìndpeaks() with a 7.5% prominence threshold. Events in separate ROIs occurring within 250 ms of each other were considered parts of the same event. (2) we reconstructed pixels and frames encompassing each event at full resolution and repeated findpeaks() using the same threshold on each pixel’s ΔF/F values.

We measured event peak intensity (Fig. 7G and 7H) from images in which each pixel was assigned its ΔF/F peak prominence value. After smoothing with a 2D gaussian filter (sigma=4) we identified 2D peaks ^40^ and used the highest value.

### Clustering

We performed hierarchical clustering on the CCF area average peak ΔF/F vectors for both IED and non-IED events. In calculating these vectors, pixels where no event peaks were detected were set to zero. All events, from both genotypes, were pooled before calculating a cluster tree with the Matlab built-in function linkage() based on the average correlation distance between clusters. To decide how many clusters to consider, we used a silhouette analysis. In the case of IEDs, a 2-cluster solution produced the highest silhouette value, so the highest local maximum was used, which was found at 9 clusters. In the case of non-IED events, a global maximum was identified at 20 clusters.

### Statistical analyses

The permutation test we used to establish significance of results in Fig. 3 is described in the figure legend. The confidence intervals used to establish significance of results in Fig. 4 are described above in the section ‘Pre-seizure image template matching’. To test for group differences in interictal activity event rate and duration (Fig. 7C, 7D & 7I), we shuffled and redrew session event rates or durations for each cluster, 10,000 times. We then calculated a *p*-value as the fraction of shuffles generating a group difference more extreme than that seen in the true arrangement. These *p*-values, as well as those in Fig. 5 were corrected for multiple comparisons using the Holm-Bonferroni method^41^ (*n* = 10 tests). Similarly, to test for event intensity differences between each individual *Kcnt1^m/m^* mouse and the WT group (Fig. 7G), we shuffled and resampled event peak intensities 10,000 times, and used Holm-Bonferroni to correct for multiple comparisons (*n* = 6 tests). To test for peak intensity differences between events in each cluster, in each *Kcnt1^m/m^* mouse and the WT group, we shuffled and resampled event intensities 10,000 times and corrected for multiple comparisons using the Bonferroni method (*n* = 60 tests, 10 clusters in 6 *Kcnt1^m/m^* mice). The significance of results in Fig. 7H are described in the figure legend.

### Data availability

The data and code supporting the findings of this study are available upon reasonable request from the corresponding author.

## Results

### Widefield imaging of the dorsal cortex in *Kcnt1^m/m^* and WT mice

To understand how a *KCNT1* GOF variant alters brain activity at the macroscopic scale, we performed widefield Ca^2+^ imaging of the dorsal cortex in *Kcnt1^m/m^* and WT mice (Fig. 1A and 1B). We measured seizures, IEDs, and interictal cortical activity (all activity outside of seizures and IEDs in both *Kcnt1^m/m^* and WT mice). Experimental mice were generated by crossing a line carrying the Y777H *KCNT1* variant (corresponding to human Y796H) to the *Snap25*-2A-GCaMP6s knockin line, in which GCaMP6s is expressed in all neurons (Fig. 1B).^33^ To gain optical access to the cortex, we either removed the dorsal skull, implanting a glass cranial window (*n* = 1 *Kcnt1^m/m^*. or rendered the bone semi-transparent with a layer of cyanoacrylate (*n* = 4 WT, *n* = 5 Kcnt1^m/m^).^17,34^ We then imaged cortical GCaMP6s fluorescence with a custom tandem-lens epifluorescent macroscope ^18,36^ while mice were awake and free to run on a Styrofoam treadwheel (Fig. 1A, Experimental Methods). In addition to cortical imaging, we recorded treadwheel motion and whole mouse video during each session. In total, we collected 34.5 hr of imaging data, 21 hr from six *Kcnt1^m/m^* mice and 13.5 hr from four WT mice.

### Spontaneous seizure mapping identified seizure susceptible cortical areas in *Kcnt1^m/m^* mice

Previously, using video-EEG monitoring, we showed that *Kcnt1^m/m^* mice have two types of spontaneous seizures: generalized tonic-clonic (lasting 30-60 sec), and tonic (≈ 5 sec, often occurred in trains). ^17^ In our widefield imaging data, we identified seizures as abnormally high GCaMP6s fluorescence occurring simultaneously with behavioral seizure correlates, such as Straub tail, back arching, and convulsions (Fig. 1C and Supplementary video 1). Of the 52 seizures we observed in four *Kcnt1^m/m^* mice, most were brief (median duration 4.525 sec) and occurred in trains (median inter-seizure interval 13.825 sec, Fig. 1D), suggesting that they corresponded to the tonic seizures identified by video-EEG.

To map seizure emergence and propagation, we determined the time at which signals on each pixel first crossed seizure threshold (see Experimental Methods). We then defined the seizure “emergence zone” as the group of pixels above threshold in the first frame of each seizure (Fig. 2A and 2B). These areas ranged in size from 0.0026 to 0.93 mm^2^ (mean 0.28 mm^2^ ± 0.0287 s.e.m), were mostly (65%) bilaterally symmetrical (Fig. 2A) and only rarely (17%) involved discontinuous patches, separated by more than 300 μm in the same hemisphere.

We localized emergence zones to cortical areas by aligning our images to the Allen Mouse Common Coordinate Framework v3 (CCF, Supplementary Fig. 1) and measuring the fraction of each emergence zone in each area in our field of view (FOV) (Fig. 2A, 2B, and 2C).^42^ Seizures most often emerged in a region where the anterior visual (VISa, a constituent of posterior parietal cortex, PTLp), retrosplenial (RSP), and anteromedial visual (VISam) area borders converge, which we call the posterior emergence zone (PEZ). This location is illustrated by the example emergence zone in Fig. 2A, peak values in the Fig. 2B heatmap, and as seizures 1-13 in the Fig. 2C heatmap. The second most common emergence zone was medial secondary motor cortex (MOs), as illustrated in seizures 36-52 in Fig. 2C.

Following emergence, seizures expanded in the cortex following a roughly sigmoidal growth curve (Fig. 2D), reaching maximum sizes of ~9-24 mm^2^ (upper limit reflects the size of our FOV). Growth rate was variable across seizures and some of this variation was systematic; seizures that emerged shortly after another seizure ended grew much more rapidly than others.

To visualize seizure propagation, we ranked the time at which pixels entered each seizure and created maps of these ranks (Fig. 2E and 2F). These showed that seizures followed a predictable pattern in their spread; the earliest recruited areas were those that served as emergence zones during other seizures. This is illustrated by the example emergence and recruitment rank maps (Fig. 2A and 2E); following emergence at the PEZ, distant medial MOs was recruited before much closer areas such as the primary visual (VISp) and barrel (SSp-bfd) cortices. Collectively, the areas prone to both seizure emergence and early recruitment included medial MOs, lateral RSP, PTLp (comprised of VISa and rostrolateral visual area, VISrl), VISam, SSp-tr, and the most posteromedial corner of MOp, which links the anterior and posterior portions of this set (Fig. 2F). These areas also showed the most intense seizure-related activity (Fig. 2G) and the longest seizure durations (Fig. 2H). Taken together, these data argue that not all cortical areas in *Kcnt1^m/m^* mice are equally susceptible to seizures; a consistent subset is most likely to host emergence, get recruited early, and show the most intense and longest duration activity during seizures.

The bias in seizure susceptibility toward specific cortical areas in *Kcnt1^m/m^* mice could be KCNT1-dependent, reflecting some feature of its expression or function, or KCNT1-independent, reflecting some intrinsic susceptibility of certain cortical regions to epileptiform activity. To distinguish between these possibilities, we compared *Kcnt1^m/m^* seizure maps to equivalent maps of seizures induced in WT mice by systemic administration of the GABA_A_ receptor antagonist picrotoxin (Supplemental Fig. 2). The cortical area most susceptible to picrotoxin seizures was MOs, which is also susceptible to *Kcnt1^m/m^* seizures and IEDs, suggesting this area is intrinsically susceptible to epileptic activity. In contrast, the PEZ was not notably susceptible to picrotoxin seizures, leading us to conclude that the pattern of seizure susceptibility in *Kcnt1^m/m^* mice likely results from both KCNT1-independent and -dependent factors.

### Normal cortico-cortical synaptic connectivity predicts long-range seizure jumps in *Kcnt1^m/m^* mice

Recent widefield Ca^2+^ imaging of seizures induced by focal chemoconvulsant application showed that epileptic activity propagates along normal synaptic connections between brain areas.^27^ However, whether healthy patterns of synaptic connectivity similarly shape spontaneous seizure topography in a chronically epileptic brain is unknown. We investigated this by comparing *Kcnt1^m/m^* seizure propagation patterns to a normative model of cortico-cortical connectivity.

Interictal cortical activity shows bilateral symmetry at the macroscopic scale, which depends, in part, on callosal connections ^43,44^ (but see ^45^). The *Kcnt1^m/m^* seizures we observed were also bilaterally symmetrical (Fig. 1 and 2), suggesting that callosal connections also supports seizure activity. Within each hemisphere, *Kcnt1^mm^* seizures, like those induced by chemoconvulsants^27,28^, propagated both contiguously and non-contiguously in our FOV.

During contiguous spread, seizures expanded outward into adjacent tissue, potentially either via high local synaptic connectivity or non-synaptic mechanisms. Non-contiguous spread involved seizure activity developing secondarily in a distant patch of cortex (Fig. 3A). We tested whether these long-range jumps occur between synaptically connected areas.

For each jump, we recorded the CCF identities of all areas in the primary seizure location and the first area recruited at the secondary location, reasoning that all primary-to-secondary area pairs represent potential pathways for seizure spread (Fig. 3B). To minimize the combinatorial complexity of the analyses, we included only the first jump from seizures in which more than one occurred. Primary seizure locations largely overlapped with seizure emergence zones, particularly the PEZ (Fig. 2B and 2C), and secondary locations most often emerged in MOs (Fig. 3C). Several seizures displayed this pattern in reverse, starting in MOs and jumping to PTLp (VISa and VISrl, combined here for correspondence to model described below).

We then determined the fraction of primary-to-secondary location area pairs that were connected in an inter-region connectivity model based on the Allen Connectivity Atlas.^1^ We compared this value to a distribution of simulated fractions after randomly repositioning primary and secondary locations. Across all jumps in our data set, the average fraction of connected areas was 0.86, higher than any observed after random repositioning (10000 repetitions, Fig. 3E). We similarly tested whether connection strength between primary-to-secondary area pairs was greater than expected by chance. The model predicted a true grand average connection strength of 0.39, higher than any we observed after randomly re-drawing strengths from the model (10000 repetitions, Fig. 3F), likely because true primary-to-secondary area pairs contains far fewer weak connections than the model at large. These analyses showed that seizure activity jumps between synaptically connected cortical areas at much greater than chance frequency and that the connections are, on average, unusually strong, arguing that normative patterns of cortico-cortical synaptic connectivity shape spontaneous seizure propagation in a genetic model.

However, connectivity and connection strength alone did not determine where seizures propagated. For example, the model predicts that VISp receives strong input from PTLp, but it was remarkably resistant to *Kcnt1^m/m^* seizures (Fig. 2, 3C, and 3D). Additionally, *Kcnt1^m/m^* seizures that began in MOs jumped most often to PTLp, even though several other regions in our FOV receive stronger inputs (Fig. 3C and 3D). Rather, secondary seizure sites appeared to form in places that both receive strong inputs from the primary location and are, themselves, seizure susceptible.

### Abnormally persistent activity in the seizure emergence zone precedes seizure onset in *Kcnt1^m/m^* mice

In the lead-up to seizure, we observed increasing activity not only in the impending emergence zone but across our FOV, indicating that seizures emerge from periods of widespread cortical activation. We investigated whether any features of this activity heralded the oncoming seizure, reasoning that this may reveal clues to the cause of seizure initiation and important potential therapeutic targets.

We first asked whether this widespread activation was abnormal in its overall intensity, reasoning that unusually strong input to the seizure emergence zone from multiple upstream regions at such times could drive it into seizure. We tested this by taking the sum of GCaMP6s ΔF/F values in each frame during the lead up to seizure and expressing these values as a percentile of all equivalent frame sums during interictal activity. While the average pre-seizure frame sum exceeded the 50^th^ percentile ~4 sec before seizure onset and climbed steadily from then on, it did not exceed the 95th percentile until 125 ms before seizure onset (Fig. 4A). This indicates that for most of its duration, the widespread activity preceding seizure does not represent an abnormal total level of cortical activity.

Another possibility is that abnormal patterns of co-active areas drive seizure emergence. To investigate this, we first defined pre-seizure activity patterns using the last ΔF/F frame prior to seizure emergence (Fig. 4B). We then generated a peak activity template for each pattern (see examples in Fig. 4D) and compared these to equivalent templates derived from all other frames in our data set. We reasoned that if pre-seizure activity is abnormally distributed, we would find few frames outside of the pre-seizure period that produce similar templates. To quantify the similarity between templates, we defined a ‘match score’ (0-1.0), calculated as the number of overlapping peak pixels divided by the total number of peak pixels (see Experimental Methods). Next, we identified the highest match score for each pre-seizure pattern (single best match), and the average scores across the top 10, 25, 50, and 100 matches (Fig. 4C) among frames free of seizures and IEDs. The single best match scores ranged from 0.52 to 0.94, with a median of 0.88. The average score distributions closely resembled the single best match score distribution, except shifted progressively more leftward (from top 10 to 100). This indicates that peak activity in the moments before many seizures emerge is not distributed abnormally, and that this finding is robust to the exact number of top matches considered.

While the spatial pattern of activity preceding *Kcnt1^m/m^* seizures is not unusual, we wondered whether the temporal dynamics of activity in this period might be. Watching ΔF/F videos, we noticed that in the lead up to seizure, cortical activity frequently assumed, and often persisted in, the pre-seizure spatial pattern. To quantify this observation, we measured the match score between each pre-seizure template and templates from frames in the prior 20 sec (Fig. 4D). These traces showed that the lead up to seizure was characterized by a ramping up of the match score. Because pre-seizure templates encompass emergence zones (data not shown), this means a progressively larger fraction of cortical activity in the pre-seizure period localized to the future site of seizure emergence. To determine whether these dynamics were specific to the pre-seizure period, we compared the average match score preceding seizure to a null distribution of average scores preceding interictal matches (see Experimental Methods, Fig. 4E). This showed that, starting 9.4 sec before seizure onset, cortical activity is significantly more concentrated in the impending seizure emergence zone than would be expected at a corresponding time point in the lead up to similar activity outside of seizures and IEDs (Fig 4F). This is true even before the pre-seizure activity level exceeds that of interictal matches (Fig. 4G period between −9.4 and −2.4 sec). Together, these analyses indicate that the preseizure period in *Kcnt1^m/m^* cortices is distinguished by an abnormal, persistent concentration of activity in the impending seizure emergence zone.

### Cortical areas susceptible to seizures and IEDs show only partial overlap in *Kcnt1^m/m^* mice

We previously showed that *Kcnt1^m/m^* mice experience brief cortical interictal epileptiform discharges (IEDs) with peak activity, on average, concentrated in MOs. ^17^ Here, we showed that MOs was also a seizure-susceptible region, suggesting that multiple disease-related abnormalities may co-localize in the *Kcnt1^m/m^* cortex. However, the precise correspondence between IED and seizure susceptibility in *Kcnt1^m/m^* cortex remains unknown.

In widefield Ca^2+^ imaging data, IEDs are distinguished by their abnormally high intensity and brief duration.^17,27,46^ To map IED susceptibility in greater detail, we first identified events in *Kcnt1^m/m^* and WT cortical activity during seizure-free, stationary periods by detecting ΔF/F peaks that exceeded a prominence threshold of 7.5% (see Experimental Methods). All brief events (<0.875 sec mean width at half peak) with unusually high peak intensity (>99^th^ percentile for WT event prominence) were categorized as IEDs. We detected many IEDs in *Kcnt1^m/m^* mice that appeared in a scatter plot of event intensity versus duration as a lobe of points not seen in WT mice (Fig. 5A and 5B, magenta points). To localize *Kcnt1^m/m^* IEDs, we measured the average peak ΔF/F in each cortical area we imaged for every IED and grouped IEDs with similar spatial profiles using hierarchical clustering (Fig. 5C, see Experimental Methods). Most *Kcnt1^m/m^* IEDs had peak activity in MOs; however, they also variably involved lower levels of activation in primary motor cortex (MOp), adjacent primary somatosensory (SSp) regions, and PTLp (1^st^, 3^rd^, and 4^th^ largest clusters in Fig. 5C). Another large population of IEDs showed peak activity in posterior retrosplenial cortex (RSP), extending into adjacent higher visual areas and posterior MOs (2^nd^ largest cluster in Fig. 5C). Together, these four clusters represent almost 90% of IEDs.

These data demonstrated that, in *Kcnt1^m/m^* cortex, the most susceptible region to IEDs is MOs, a region also highly susceptible to seizure; however, IEDs with peak activity in MOs tended not to involve other seizure-susceptible regions, but rather SSp areas that rarely participate in seizures. This could be a result of seizures and IEDs engaging distinct populations of MOs neurons with different connectivity or, less likely in our minds, a different set of rules governing propagation of these events. Moreover, apart from MOs, we found that seizure and IED-susceptible regions were largely non-overlapping in the *Kcnt1^m/m^* cortex. For instance, although RSP was prone to both, the sub-regions involved in each appeared distinct (Fig. 2B and 5C). Importantly, we did not find any IED cluster with peak activity at the most frequent seizure emergence site, the PEZ (Fig. 2B and 5C), suggesting that the neural population most likely to host seizure emergence is not notably susceptible to IEDs.

### IEDs constitute the high intensity tail of excessive interictal activity centered in *Kcnt1^m/m^* MOs

We classify IEDs as pathological because brief events of equivalent intensity are exceedingly rare in WT mice. However, IEDs do not appear as a distinct cluster of points in the *Kcnt1^m/m^* intensity versus duration plot, but rather a continuous extension of the point cloud (Fig. 5A). This suggests that there may not be a clear distinction between IEDs and non-IED events, and that the mechanism generating IEDs may also produce activity outside of our IED thresholds.

To investigate, we first measured the rate and spatial distribution of non-IED events in *Kcnt1^m/m^* and WT mice. Reasoning that the characteristic short duration of IEDs might also be seen in pathologically driven non-IED events, we split these into two groups: brief, low intensity events (Fig. 5A and 5B, blue dots) and all prolonged events (Fig. 5A and 5B, black dots). We then calculated the rate of occurrence of both groups at every pixel (Fig. 5D and 5E). Strikingly, the rate of brief events in IED susceptible areas (MOs, MOp, and adjacent SSp) was much higher in *Kcnt1^m/m^* than WT mice, whereas the rate and spatial distribution of prolonged events was similar across genotypes. This demonstrates that *Kcnt1^m/m^* mice suffer from excessive cortical activity centered in MOs, and that IEDs are simply the highest intensity events associated with this abnormality. More generally, it shows that the *Kcnt1* variant affects brain activity outside of seizure and IED, and suggests that these dramatic events may be extreme, and relatively rare, outcomes of underlying defects that mostly impact interictal activity.

### Spatial organization of interictal cortical activity is largely preserved in the *Kcnt1^m/m^* mice

The difference in brief event rate maps we observed across genotypes (Fig. 5D and 5E) might reflect a novel event type in *Kcnt1^m/m^* cortex, in which MOs is abnormally activated alongside SSp areas only in this genotype. To investigate whether the spatial organization of interictal cortical activity is disrupted in *Kcnt1^m/m^* mice, we considered all events detected outside of seizure, IED, and treadmill movement (event detection described above, see Experimental Methods). Similar to our IED analysis, we measured the average peak intensity of neural activity within each cortical area during every event and grouped events with similar spatial profiles using hierarchical clustering. Silhouette analysis identified a natural grouping in the data consisting of 20 clusters. The 10 largest of these contained 99.5% of events and were our exclusive focus in the analyses that follow. For each cluster, we calculated the average peak intensity of neural activity in each cortical area across all events in the cluster, first within individual mice and then across *Kcnt1^m/m^* and WT groups (Fig. 6, line plots). Additionally, we generated average event peak intensity images for each cluster and genotype (Fig. 6, images).

Two features of these data indicate that events in each cluster represent event types that occur commonly, and are equivalent, in both WT and *Kcnt1^m/m^* mice. First, even though events from both groups were first pooled and clustering performed without respect to genotype, each cluster contained many events from both WT and *Kcnt1^m/m^* mice (Fig. 6, line plot text insets).

If the spatial organization of events in *Kcnt1^m/m^* mice differed significantly from WT, we would expect clusters comprised mainly of one genotype or the other. Secondly, events from both groups in the same cluster showed remarkably similar spatial profiles, as evidenced by their average peak intensity images and by the high positive correlation coefficient between the peak intensity traces (significant at 0.005 for all clusters, median = 0.97, std = 0.05). The clustering algorithm attempts to maximize within-cluster correlations by design, but because events were pooled across groups before clustering, the degree to which it found similar events from both groups represents a biological result. This does not mean events in the same cluster were identical in WT and *Kcnt1^m/m^* mice; in some clusters, we found group differences in average peak intensity in some cortical areas. However, even in these cases, the spatial distribution of activity was highly correlated across groups. These data led us to conclude that the major types of cortical activity observed in WT mice are preserved in *Kcnt1^m/m^* mice. Also, we found no evidence of any major novel event types in *Kcnt1^m/m^* mice during interictal cortical activity, suggesting that large scale disruption of cortical arealization and connectivity are not necessary to support frequent spontaneous seizures and IEDs and further, that seizures and IEDs do not necessarily cause such disruptions.

### Excessive interictal activity in *Kcnt1^m/m^* cortical areas and mice prone to seizure and IED

Because novel associations of cortical areas during interictal activity appear rare in *Kcnt1^m/m^* mice (Fig. 6), we reasoned that the difference in brief event rate maps we observed (Fig. 5D and 5E) likely reflects a change in the rate and/or intensity of a common event type. To identify such abnormalities in *Kcnt1^m/m^* interictal activity, we compared three parameters of each event cluster across genotypes: event rate, peak intensity, and duration.

We measured the WT and *Kcnt1^m/m^* event rate for each cluster in all sessions where event detection was performed. Because the number of sessions per animal differed, we calculated group averages weighted by the number of sessions from each mouse (Fig. 7A and 7B). Using a permutation test to establish significance, we found group event rate differences in 5 out of the 10 clusters (Fig. 7C-7F). Four showed increased rate in *Kcnt1^m/m^* mice, and one decreased. Those increased in *Kcnt1^m/m^* mice had peak activity in cortical areas susceptible to seizures and IEDs (Fig. 7C, 7E, and 7F), whereas the cluster that showed a decrease localized to SSp-ul, an area not notably susceptible to seizures or IEDs (Fig. 7D). These results suggest that a distinguishing feature of tissue prone to epileptic pathology is the presence of excessively frequent events during interictal periods.

Motivated by the observation that interictal activity in MOs appeared more intense in *Kcnt1^m/m^* mice (Fig. 5), we next investigated whether this holds true for all seizure and IED prone areas. Surprisingly, when we measured the peak intensity of events in each cluster, we found greater variation across mice than cluster in the *Kcnt1^m/m^*, but not WT, group. Some *Kcnt1^m/m^* mice produced events with characteristically lower intensity than WT, and others higher. An exception was the cluster centered on MOs. These events were more intense in all *Kcnt1^m/m^* mice, except the individual with the lowest average event intensity (see Experimental Methods, data not shown). To test this intensity effect at the level of individual mice, we pooled events from all clusters in each *Kcnt1^m/m^* mouse and used a permutation test to compare its session averaged intensity to the WT group. All but one *Kcnt1^m/m^* mouse differed significantly from WT (Fig. 7G). Interestingly, two of the three mice showing reduced event intensity did not experience any seizures during imaging, while the two with increased event intensity had the highest cumulative time-in-seizure. To test whether the intensity of interictal cortical activity predicted a mouse’s seizure burden, we plotted total time-in-seizure against session averaged event intensity and found these values were significantly correlated (Fig. 7H, upper plot, Spearman’s rank correlation ρ = 0.93, *p* = 0.02). We also tested whether event intensity predicted IED rate and found a positive correlation that was not quite significant at α = 0.05 (Fig. 7H, lower plot, Spearman’s rank correlation, p = 0.83, *p* = 0.06). Together, these results indicate that the YH variant exerts a bidirectional effect on the intensity of interictal cortical activity and, in contrast to event rate, the strongest determinant was not event type but individual. Most strikingly, the intensity of interictal cortical activity predicted a mouse’s seizure burden (Fig. 7H).

Because duration is a distinguishing feature of both pre-seizure activity and IEDs, we next investigated whether this parameter was altered in interictal *Kcnt1^m/m^* cortical activity. To compare event duration between genotypes, we again calculated group averages for each cluster and tested differences with a permutation test (see Experimental Methods). Similar to our observations with event rate and intensity, we found significant duration differences in both directions; in eight clusters, the duration of *Kcnt1^m/m^* events was significantly reduced, and in two, it was significantly increased (Fig. 7I). The largest reduction was among events centered on MOs, which also occurred at a higher rate and intensity in *Kcnt1^m/m^* mice (Fig. 7C, Fig. 5).

The next three largest reductions were also in clusters centered on seizure- and IED-susceptible areas (Fig 7I, asterisks). These data suggest that reduced duration might be linked to increased intensity and rate, either due to some shared underlying cause or as a compensatory change triggered by excess activity such as increased feedback inhibition.

Taken together these results show that, outside of seizures and IEDs, the YH variant bidirectionally modulates the rate, intensity, and duration of cortical activity, depending on area and individual. Critically, seizures and IEDs were specifically associated with areas and mice where interictal cortical activity was excessive. The scope and diversity of interictal effects coupled with this specific association emphasizes the need to establish links between any given variant effect and disease symptoms to efficiently target future research and therapies.

## Discussion

Spontaneous seizures are a defining feature of epileptic disease. Yet how the root cause of seizures impacts interictal brain activity is poorly understood. This is an important gap in our understanding for several reasons. Subtle abnormalities in the epileptic brain activity may (1) lead to seizure initiation, either directly or indirectly, (2) identify seizure initiation zones and (3) underlie comorbidities. In each scenario, such abnormalities represent valuable potential therapeutic targets.

Part of the reason subtle differences between healthy and epileptic brains have remained mysterious is methodological. The most common method used to study spontaneous seizures is EEG recording.^47^ While this technique offers spectacular temporal resolution and can sample large areas of tissue with electrode arrays, a drawback is that the relationship between underlying neural activity and the EEG signal is not straightforward. Currents detectable on the EEG may be caused by either neuronal firing, synaptic activity, or other ion fluxes. Large, synchronous neural activations, as in seizure and interictal spiking, are clear in EEG traces, but less intense and/or synchronous events, which comprise most brain activity, are more difficult to detect and study. Widefield Ca^2+^ imaging provides a more linear read-out of neural activation and is sensitive enough to reveal normal spontaneous and sensory-evoked activity. Using this technique allowed us to image seizure and IED activity, as well as interictal cortical activity, in a genetic mouse model of human disease. Critically, we also collected equivalent data from WT mice, allowing us to define ‘normal’ in a non-epileptic brain. From these data, we made two valuable comparisons: seizures and IEDs compared to interictal activity in *Kcnt1^m/m^* mice, and interictal activity in *Kcnt1^m/m^* mice compared to WT mice. This process demonstrated that an epilepsy-causing gene variant has strong, widespread, and cortical region-specific effects on brain activity outside of seizure and IEDs, which builds upon prior findings in a genetic zebrafish model. ^48^ Importantly, while interictal activity in epileptic animals differed in numerous ways from WT, not all were associated with seizures and IEDs; epileptic activity occurred where interictal activity was excessive.

Previously, we showed that a major effect of the YH variant on cellular physiology is a reduction of the intrinsic excitability of inhibitory, but not excitatory, neurons from motor cortex. ^17^ While this offers a potential explanation for increased event rate and/or intensity, what underlies reductions in these parameters is more mysterious. Perhaps, reduced excitability is not specific to inhibitory neurons in other cortical areas and/or upstream brain regions. The variation across *Kcnt1^m/m^* mice in event intensity might reflect differences in disease progression or a strong dependence of disease outcome on individual biology. It is interesting to note that the Na^+^-activated K^+^ current, to which KCNT1 channels contribute, has long been proposed to mediate a slow afterhyperpolarization in neurons following periods of activation.^49–51^ Increasing the magnitude of such a current could reduce the duration of activity, offering a potential explanation for the widespread reduction in event duration we observed. Clearly, many details beyond changes in channel function and cellular physiology are needed to explain YH variant effects at the macroscopic scale. How the role of KCNT1 channels differs across brain areas and development, as well as what indirect effects and compensatory processes the variant triggers, are all important questions for future study.

One limitation of widefield imaging is that, while we monitored a large fraction of the cortex with high spatial resolution, we were unable to monitor activity in the entire brain, including several subcortical structures known to play important role in seizure generation. This raises the concern that we did not image the brain areas responsible for generating pathological activity, only its propagation into our FOV. Although it is almost certain that areas outside our FOV are playing a role, several aspects of our data are inconsistent with exclusively remote seizure and IED initiation. Our core finding of excessive interictal activity in susceptible areas argues that they are at least a portion of the generative network. Such correlations in space and time between interictal abnormalities and epileptiform activity are hard to reconcile with a purely remote seizure and IED origin. For example, abnormally persistent activity in the seizure emergence zone begins over 9 sec before seizure onset and becomes more pronounced as seizure approaches. During this period, we are unable to see any behavioral indication of seizure, although these signs commence simultaneously with cortical seizure onset.

The high frequency of spontaneous seizures in our *Kcnt1^m/m^* mice presented a unique opportunity to image these events at the macroscopic scale. Previous seizure studies using this approach have relied on chemoconvulsants or introduction of tumor cells, and a comparison to these models is informative.^27,28^ Our finding that seizure discontinuities span synaptically connected areas is in agreement with the conclusion of Rossi et al.^27^ that seizures resulting from local application of picrotoxin and pilocarpine propagate along normal underlying synaptic pathways. Additionally, the fact that discontinuities linked seizure susceptible areas is similar to the findings in Liou et al.^28^ that long-range seizure jumps occur preferentially to distant, pharmacologically disinhibited areas. This parallel suggests that seizure susceptibility in *Kcnt1^m/m^* mice may coincide with areas with the most severely compromised inhibition. However, an important difference between these models is that chemoconvulsants fail to reveal the set of pathologically susceptibility cortical areas seen in *Kcnt1^m/m^* mice (Supplementary Fig. 2). This is important because not all areas are affected in the same way by *Kcnt1* mutation, and knowing where abnormalities occur and how they relate to disease symptoms is essential for targeting research and ultimately therapies. Furthermore, because of the strong effect chemoconvulsant drugs have on brain activity, even outside of seizures, it’s uncertain whether they can be used to relate features of interictal and pre-ictal activity to seizure occurrence in a way that is relevant to chronic epilepsy.

In this study, we show that widefield Ca^2+^ imaging can be applied to identify cortical networks underlying disease in mouse models of epilepsy. This is especially needed when the localization of pathology is not known *a priori*, as in the majority of genetic mouse models carrying human disease-causing variants. Furthermore, we were able to identify correlations between abnormalities in interictal activity and symptom-related pathological activity, a critical step to linking dysfunction across scales. Demonstrating that seizure and IEDs occur in areas that also show excessive interictal activity opens the door to investigate how underlying cellular and synaptic defects give rise to excess activity and whether interventions that target them may reduce seizure burden.

## Abbreviations

ADNFLE: autosomal dominant nocturnal frontal lobe epilepsy
FOV: field of view
CCF: common coordinate framework
CSD: cortical spreading depression
GOF: gain-of-function
IED: interictal epileptiform discharge
PEZ: posterior emergence zone
SPCM: seed pixel correlation map
SVD: singular value decomposition
SHE: sleep-related hypermotor epilepsy
WT: wild type
YH: Y777H Kcnt1 variant

## Acknowledgements

We would like to thank Todd Clason for his assistance with imaging equipment, Dr. Matt Mahoney for assistance with statistics, data analysis and coding, Dr. Amy Shore for help with genotyping and manuscript editing, the Vermont Advanced Computing Center, members of the Weston lab and ECD group at UVM for their feedback and Wayne Frankel and his group at Colombia for the Kcnt1 Y777H mouse line.

---

**Supplementary Figure S1.**
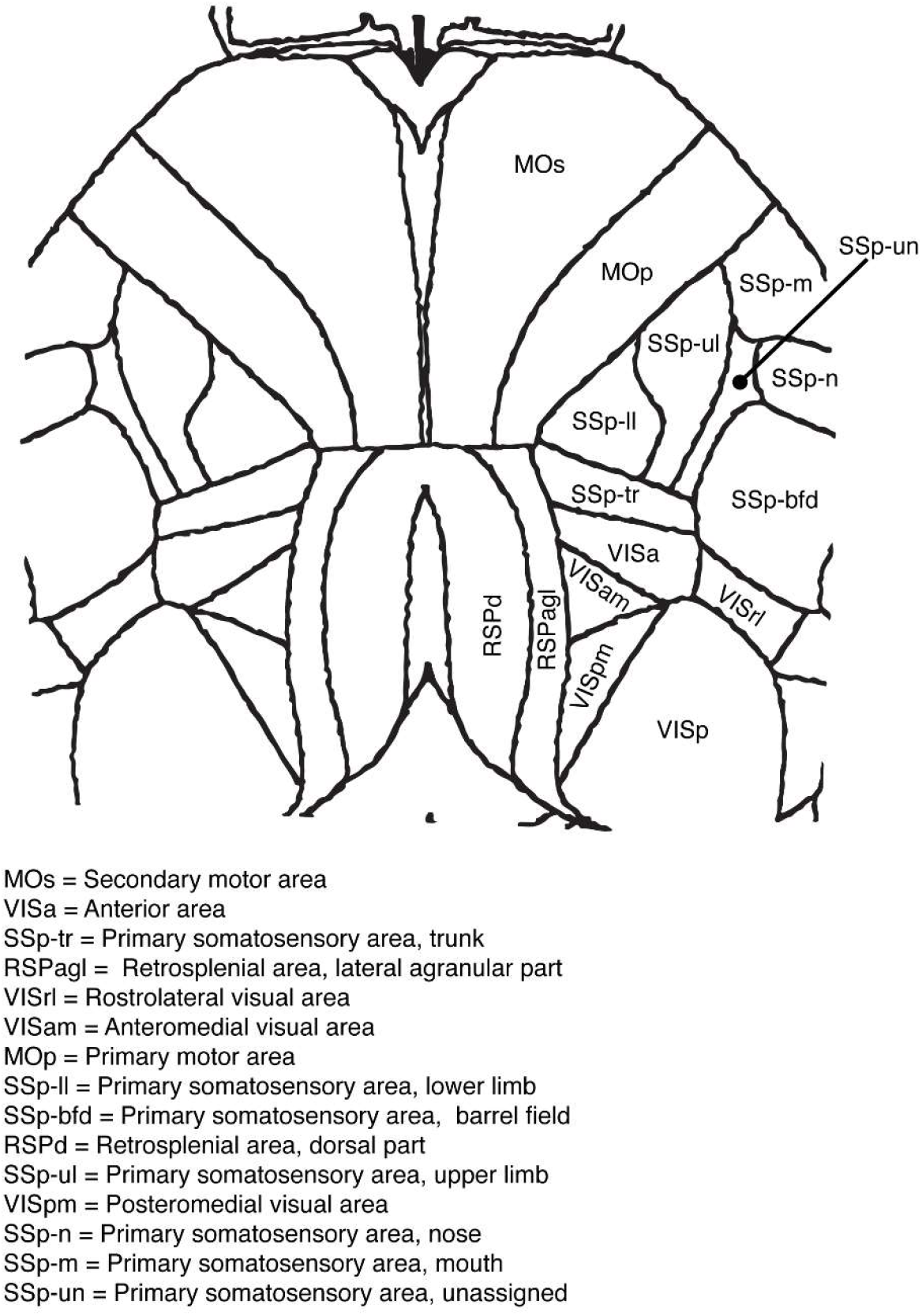
Allen Mouse Common Coordinate Framework (CCF) dorsal cortex map with labels; includes areas imaged in all mice and sessions

**Supplementary Figure S2.**
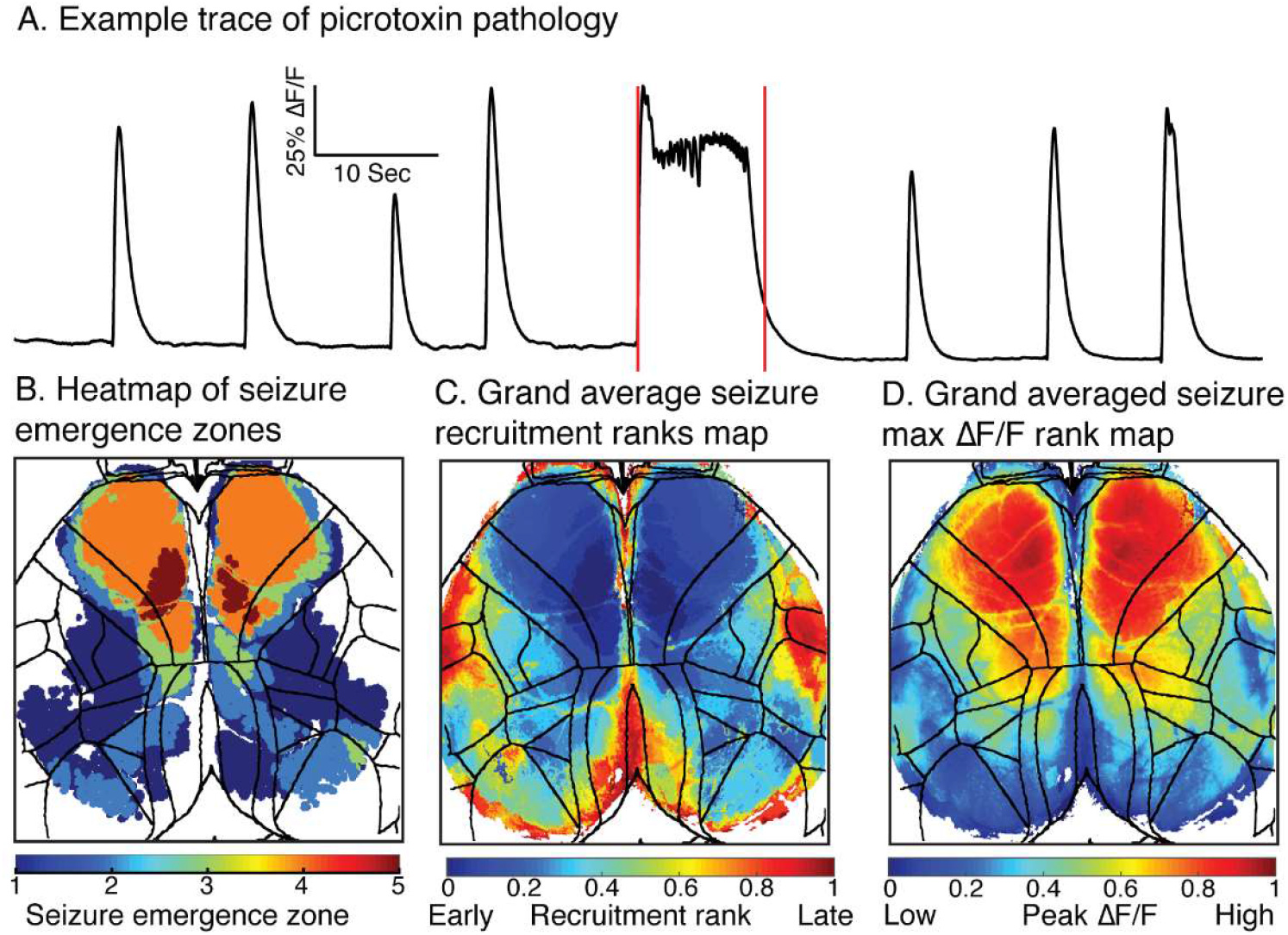
Picrotoxin seizures in WT mice Picrotoxin seizures in WT mice highlight the intrinsic susceptibility of MOs to epileptic activity. (A) An example average ΔF/F trace displaying several brief epileptiform discharges and a seizure (demarcated by red lines) induced by systemic picrotoxin administration. (B) A heatmap of seizure emergence zones summed over all five picrotoxin seizures. The value of each pixel indicates the number of seizures in which the pixel was a part of the emergence zone. (C) A grand average seizure recruitment rank heatmap, taken across all individual mouse average maps of picrotoxin seizures after co-registration. (D) A grand average heatmap of ranked pixel peak ΔF/F during picrotoxin seizures, taken across individual mouse averages.

